# Principles of nutrition-dependent root-microbiome engineering for regulating crop yield via live microbial inoculants

**DOI:** 10.1101/2024.10.25.620170

**Authors:** Yuxiao Huang, Shengyue Tang, Rumeng Liu, Ju Liu, Ting Xiao, Bin Ni

## Abstract

As the second genome, host-associated microbiota can enhance the functions of host immune systems, improve host nutrient availability and acquisition capacity, and increase host resistance to abiotic stress. Constructing host-associated microbiomes by applying various bioinoculants has become an attractive tool for promoting human health and increasing food production. However, despite their promising properties, the efficacy of bioinoculants vary significantly in actual field and clinical practices. Understanding the global design principles that shape the outcomes of interactions between bioinoculants and target host–microbiome symbionts remains a considerable challenge. In this study, we used a wheat production system as a case study to quantitatively understand how soil nutrient status impacts the establishment of host–microbiome interaction networks and their subsequent interactions with external bioinoculants. We found that soil organic carbon, one of the most general soil properties for global crop productivity and resilience, could affect the outcome of bioinoculant applications in wheat production systems, which led to functional instability in bioinoculant application outcomes. The results of this study significantly improved our understanding of the global design principles of nutrition-dependent root-microbiome engineering for regulating crop yield via the application of live microbial inoculants and provide theoretical guidance for bioinoculant applications in agricultural practice.

## Introduction

Global crop demand is growing and will likely continue to increase for decades due to increases in the world population and per capita consumption. However, crop yields are predicted to increase to meet the projected demand in 2050^1^. Due to the intensive use of chemical fertilizers, pesticides, climate change, environmental pollution, and biotic stresses from pests or pathogens, there is tremendous pressure on agricultural crop productivity^2^. It remains an enormous challenge to sustainably increase the productivity of agroecosystems. As one of the most widely cultivated cereal crops in the world, wheat (*Triticum aestivum* L.) provides ∼20% of the calories and proteins consumed by humans^3^. The contributions of wheat breeding efforts to grain yield have been significant and have played an essential role in ensuring food security for millions of people worldwide^4,5^. To improve the productivity of wheat crops, breeders have devoted significant efforts to identify and select plants with desirable traits. These traits include large seeds, high protein content, good adaptation to local environments, and resistance to pests and diseases. One of the most well-known examples of these efforts is the "green revolution" of the 1960s, which resulted in a substantial increase in cereal crop yield through the widespread cultivation of semidwarf and lodging-resistant varieties. This was achieved by manipulating plant architecture using *reduced height-B1b* (*Rht-B1b*) and *Rht-D1b* alleles^6^.

As the second genome of plants, the root-associated microbiome has largely unexplored potential to expand the genomic capabilities of host crops by providing enhanced nutrient acquisition systems, improved root architecture, and tolerance to biotic or abiotic stress^7-9^. Harnessing the regulatory power of the crop root-associated microbiome has been postulated to be a promising approach to overcome challenges related to food security and environmental health^10^. Through microbiome engineering, researchers seek to improve the functions of ecosystems by manipulating the composition and function of host-associated microbial communities to boost human health or agricultural productivity^11^. To date, two engineering strategies involving the manipulation of indigenous microbial communities or bioinoculant design and delivery have been widely used^12^. Despite their promising properties and the technical advances to date, bioinoculants fail to reshape or have long-lasting effects on microbiome functions in medicinal probiotics and microbial fertilizers in agriculture. The efficacy of these bioinoculants may vary dramatically under different environmental conditions or host genetic backgrounds^13,14^. Understanding the global design principles involved in host–microbiome assembly and bioinoculant success in complex environments such as the mammalian gut or the rhizosphere remains challenging.

The rhizosphere is a narrow region of soil that surrounds and is influenced by plant roots. It provides a multitude of niches for the growth of bacteria, fungi, viruses, *etc*., forming complex association networks with plants^15^. Microorganisms living within and on the host are essential for the overall health of the host. Moreover, nutrition plays an important role in the relationship between hosts and their associated microbiota. On the one hand, the host diet provides the nutrients necessary for the growth and maintenance of the microbiome. On the other hand, the microbiome can affect the host’s nutritional status, metabolism, and immune system^16^. For example, certain microorganisms can produce vitamins and other nutrients that the host needs, while others can compete with the host for nutrients. Nutrient concentration or quality has been reported to affect the physiological state of different microbial species, their pattern of interactions, species richness, and the stability of microbial communities^17-19^. Previous studies have also indicated that nutrition is crucial to the success of fecal microbiome transplantation (FMT). A high-fiber diet rich in fruits, vegetables, and whole grains can help to promote the growth of beneficial bacteria and reduce the risk of *Clostridioides difficile* infection. In addition, pre- and post-FMT nutrition therapy with probiotics, prebiotics, and synbiotics has been shown to positively affect the gut microbiome and the clinical outcomes of FMT^14,20,21^.

Soil organic carbon (SOC) is a complex mixture of organic compounds derived from the decomposition of plant and animal materials deposited in the soil over time. Many studies have indicated that soil organic matter is a crucial factor affecting agricultural production, nutrient availability, soil stability, and the flux of greenhouse gases^22-24^. It represents a significant carbon pool within the biosphere and is an important source and sink for carbon and nutrients. Research has shown that the presence of SOC in the soil can influence the diversity and composition of the root microbiome, with higher SOC levels typically corresponding to a more diverse and complex microbiome. This is likely because SOC provides a stable and consistent source of nutrients and energy for microorganisms and can stimulate their growth and activity. In agriculture, continuous organic fertilization strategies are widely used to regenerate soil organic carbon and enhance bacterial community diversity and microbial function to increase crop yield^25,26^.

In this study, we supplemented soil organic carbon with increasing levels of organic fertilizer to obtain a qualitative understanding of the global principles governing the effect of nutrient supply on the construction and stability of crop root–microbiome interaction networks. Additionally, effective microorganisms (EMs), one of the most widely used bioinoculants in agriculture, were employed to challenge the root microbiome of these crop root–microbiome associations, aiming to quantitatively understand the nutrition-dependent microbiome engineering principles for bioinoculants in regulating crop yield.

## Materials and methods

### Pot experiment and soil sampling

The experiments were conducted in a greenhouse at China Agricultural University (40°1′45″ N, 116°17′11″ E). The experimental soil was obtained from the Quzhou Experimental Station of China Agricultural University, Hebei Province (36°51′48″ N, 115°0′59″ E). The first experiment consisted of seven organic fertilizer treatments with six replicates in each treatment. The seven organic fertilizer treatments included no organic fertilizer (OF0), 1% organic fertilizer (OF1), 3% organic fertilizer (OF3), 5% organic fertilizer (OF5), 10% organic fertilizer (OF10), 20% organic fertilizer (OF20), and 30% organic fertilizer (OF30).

The second experiment involved combining organic fertilizers and an EM agent. The base treatment included three different levels of organic fertilizers, and within each base treatment, there were four different EM compositions with six replicates in each treatment. The three levels of organic fertilizers included no organic fertilizer (OF0), 5% organic fertilizer (OF5), and 20% organic fertilizer (O F20). Under each base treatment, there were four EM treatments: no treatment, treatment with the EM agent, treatment with the EM supernatant, and treatment with the EM pellet. The applied peach branch-based organic fertilizer contained 2.69 mg kg^-1^ NO_3_ ^-^-N, 52.03 mg kg^-1^ NH_4_^+^-N, 198.36 mg kg^-1^ AP, and 2.37 g kg^-1^ AK. Urea (46% N), superphosphate (18% P_2_O_5_), and potassium chloride (60% K_2_O) were used as the inorganic N, P, and K fertilizers, respectively.

Rhizosphere soil samples were collected at the jointing stage after 60 days of wheat growth. Wheat samples were collected for each treatment, and the roots of the soil were shaken. Soil attached to the surface of wheat roots was considered rhizosphere soil. The roots were cut off the plants with sterilized scissors and placed in sterile bags. Bulk soil samples were collected at both the jointing and maturity stages. Plant height and stem diameter were also measured during the jointing stage, while grain number and yield were recorded at the maturity stage. All soil and root samples were stored in an ice box and transported to the laboratory. The rhizosphere soil samples were collected for microbial analysis, while the bulk soil samples were used for physicochemical analyses. The soil for microbial analysis was stored at -20 ℃, while those used for determining physicochemical properties were stored at 4 ℃.

### Preparation of EM, EM pellet, and EM supernatant

EM agent is a combination of lactic bacteria, photosynthetic bacteria, yeast, and other beneficial microorganisms, assigned as EM1 from Yimu Biotechnology (Nanjing) Co., LTD in China. EM agents were centrifuged at 5000 r.p.m. at 4 ℃ for 5 minutes. The supernatant obtained by centrifugation was defined as EM supernatant, and the precipitates were washed three times in deionized water and defined as EM pellets. During all the treatments, EM/EM supernatant /EM pellet was diluted to 1:200 (v/v) and applied once a week. To keep the activity of the agents, all treatments were stored at 4 ℃.

### Cultivation and identification of rhizosphere bacteria and fungi

Rhizosphere bacteria and fungi were isolated from the greenhouse soil under the aforementioned conditions; specifically, they were isolated from the rhizosphere soil of OF0, the OF0+EM supernatant, and the OF0+EM pellet treatments. The bacteria were isolated using a high-throughput culture identification method as previously described^53,54^. In brief, the wheat roots were washed in sterile PBS buffer (0.02 M, pH 7.0) on a shaker for 5 min to collect the rhizosphere soil. The rhizosphere soil supernatants were diluted 10^-5^ times and cultivated in 96-well microtiter plates in 1/10 tryptic soy broth (TSB) for two weeks at 28 °C. A two-sided barcode Illumina PCR system was utilized for the identification of high-purity bacteria. Subsequently, nonredundant strains were screened if their 16S rDNA sequence homology was less than 97%.

We isolated and identified fungi using traditional methods to obtain a collection of fungi; this involved colony picking and Sanger sequencing. The rhizosphere soil of three samples from each treatment was pooled to increase the diversity of fungi. The rhizosphere soil was subjected to gradient dilutions of 10^-3^, 10^-4^ and 10^-5^ in PBS buffer for coating PDA plates (containing 1% Bengal red and 1% streptomycin). Fungi were purified by transplanting onto new PDA plates. Pure fungi were identified by Sanger sequencing with ITS1F and ITS4R primers and used to prepare glycerol stocks for culture collection.

### Assessment of wheat growth by rhizosphere bacteria and fungi

The impact of all the isolated bacteria and fungi on wheat growth was assessed using a seedling tray method. Each pot contained 2 wheat plants and 200 g of soil. Subsequently, 5 mL of bacterial suspension or fungal spore suspension was inoculated during the seedling stage. Bacterial suspensions were prepared through overnight culture and washed three times in sterile water, while fungal spore suspensions were prepared by washing spores in sterile water. After 3 weeks, the height of each plant was recorded, and the dry weights of the plants and roots were measured. All the isolates were tested with at least two replicates.

### Soil physicochemical property analysis

Bulk impurities were removed from the fresh soil samples. Soil pH was measured by a pH electrode at a soil:water ratio of 1:2.5. Fresh soil samples were extracted with 0.01 M CaCl_2_, and NH_4_^+^-N and NO_3_^-^-N were measured using a flow injection autoanalyzer. AP was measured using a flame spectrophotometer, and AK was determined by the Mo-Sb colorimetric method. SOC was measured using the potassium dichromate method.

### Extraction of DNA, amplicon sequencing, and bioinformatic analyses

To separate rhizosphere soil samples, root samples were placed in 50 mL sterile tubes and vigorously shaken after 40 mL of sterile phosphate-buffered saline (PBS) solution was added. The samples were then washed with PBS 5-6 times to collect the rhizosphere soil attached to the root surface. Subsequently, the tubes were centrifuged, and the pellets were collected as the rhizosphere soil sample. All the samples were stored at -20 °C for DNA extraction. Genomic DNA was extracted using the CTAB/SDS method, and its concentration and purity were monitored on 1% agarose gels. Based on the concentration, the DNA was diluted to l μg/μL using sterile water.

Prokaryotic 16S rRNA genes were amplified using primers 341F (5′-CCTAYGGGRBGCASCAG-3′) and 806R (5′-GGACTACNNGGGTATCTAAT-3′) with barcodes. ITS genes were amplified using primers ITS1F (5’-CTTGGTCATTTAGAGGAAGTAA-3’) and ITS2R (5’-GCTGCGTTCTTCATCGATGC-3’) with barcodes. All PCRs were carried out with 15 μL of Phusion® High-Fidelity PCR Master Mix (New England Biolabs), 0.2 μM forward and reverse primers, and approximately 10 ng of template DNA. Thermal cycling consisted of initial denaturation at 98 °C for 1 min, followed by 30 cycles of denaturation at 98 °C for 10 s, annealing at 50 °C for 30 s, elongation at 72 °C for 30 s, and finally 72 °C for 5 min. PCR products were mixed in equal ratios. Then, the mixed PCR products were purified with a Qiagen Gel Extraction Kit (Qiagen, Germany). Sequencing libraries were generated using a TruSeq® DNA PCR-Free Sample Preparation Kit (Illumina, USA) following the manufacturer’s recommendations, and index codes were added. Library quality was assessed on a Qubit@2.0 fluorometer (Thermo Scientific) and an Agilent Bioanalyzer 2100 system. Finally, the library was sequenced using an Illumina NovaSeq platform, which generated 250 bp paired-end reads.

Paired-end reads were merged using FLASH (VI.2.7, http://ccb.jhu.edu/software/FLASH/)^55^. Quality filtering of the raw tags was performed under specific filtering conditions to obtain high-quality clean tags according to FASTP^56^. The tags were compared with the reference database (Silva database, https://www.arb-silva.de/) using the UCHIME algorithm (UCHIME Algorithm, http://www.drive5.com/usearch/manual/uchime) to detect chimeric sequences, after which the chimeric sequences were removed^57,58^.

### Statistical analyses

Statistical differences were determined by one-way ANOVA using SPSS statistical software (version 26.0, IBM, Armonk, NY, USA). The means were compared using Duncan’s test (*p* < 0.05). The figures were made using the R software package (https://www.r-project.org/, version 4.2.1).

### Co-occurrence network construction

The bacterial and fungal association networks were constructed based on the relative abundance datasets of the OTUs. OTUs detected in less than 30% of the samples were discarded to improve the statistical power of the correlation calculations while minimizing the loss of OTUs. The correlation matrix for OTUs was calculated using Spearman’s rank-based correlation. The appropriate correlation coefficient cutoff for defining the network was determined automatically using a random matrix theory (RMT)-based approach. The network modules were detected by fast greedy modularity optimization. Additionally, random networks corresponding to each empirical network were constructed by keeping the numbers of nodes and links constant and rewiring the nodes. The topological indices for both the empirical and random networks were calculated as described using the network analysis pipeline at http://ieg4.rccc.ou.edu/mena/. Structural equation modeling was also employed in this study.

### Quantification of normalized stochasticity ratio in the rhizosphere

The normalized stochasticity ratio (NST) is a new metric that quantifies the relative importance of stochasticity in governing community assembly, ranging from 0 to 1; values higher than 0.5 indicate more stochastic assembly, while those less than 0.5 indicate more deterministic assembly. NST reflects the contribution of stochastic processes based on relative differences between observed communities and a null expectation, which was conducted using the tNST function in the “NST” package^59^.

### Structural equation modeling

To further distinguish the direct and indirect effects of environmental drivers on microbial biodiversity, we conducted structural equation modeling (SEM) analyses to investigate the relationships among the experimental treatments, soil and plant variables, and microbial diversity. To account for potential temporal autocorrelation, we utilized plot-level data by averaging the microbial or environmental data across timepoints within the same plot. Initially, we developed a conceptual model that included all reasonable pathways. Subsequently, nonsignificant pathways were sequentially eliminated unless they provided biologically informative insights, or additional pathways were added based on residual correlations. This process was repeated until the model demonstrated adequate fit, with P values of the χ2 test > 0.05 (indicating no significant difference between the predicted model and observed data) and a root mean square error of approximation (RMSE) < 0.08. SEM analysis was conducted using the lavaan R package.

### Data collection for Meta-analysis

A comprehensive scientific literature search was performed using the Web of Science database (http://apps.webofknowledge.com/) and China National Knowledge Infrastructure (CNKI) until January 2024. The keywords were ‘compound microbial fertilizer or bio-organic fertilizer or microbial inoculant’, ‘yield’, ‘soil organic carbon’. The sample criteria were as follows: 1. Pairwise comparisons were made between the application of microbial fertilizers and the control group without microbial fertilizers under the same soil climate conditions. Only studies that conducted pairwise comparisons between microbial fertilizer application and non-application under the same levels of inorganic and organic fertilizer inputs were included; 2. The experiments must include average values, standard errors or standard deviations, and repetition numbers of yield to calculate different levels of efficiency and effect sizes; 3. The studies must be field experiments, including essential information such as soil physicochemical properties and nutrient addition treatments. Based on the above screening criteria, 668 sets of data from 139 articles were collected. Data presented in tabular form in the literature was directly extracted, while data presented in other graphical forms, such as bar charts, line charts, etc., were digitized using the software Get Data Graph Digitizer 2.24.

### Global Meta-analysis

The meta-analysis was conducted with R Software Version 3.3.3 and the interface R-Studio Version using the “metafor” package. In our dataset, the observations were distributed across all geographic regions. Soil organic carbon (SOC) content was classified into three classes: low (≤17.6 g kg^-1^), moderate (17.6-35.2 g kg^-1^), and high (>35.2 g kg^-1^). The impact of SOC on yield was tested in a model of mixed-effects meta-regression using the “glmulti” package in R. The relative importance value for each prediction was equal to the sum model of the Akaike weights in which that prediction appears. These values can be viewed as the overall support for each variable across all models. A cut-off of 0.8 was set to differentiate between essential and non-essential predictors^60^. The natural log-transformed response ratio (lnR) was used to quantify the effect of microbial fertilizers on crop yield. The lnR, also called “effect size”, was dimensionless and used to characterize the relative changes between treatments and control^42^.

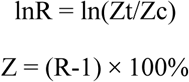

Zt and Zc are the mean values of the response variables in the treatment group (t) and control group (c). The results, reported as percentage changes (Z) under the treatment effect, were found to be significant when the 95% CIs did not overlap with zero^61^. The thoroughness of our analysis is evident in the significant results of Kendall’s tau bias test and the substantial number of fail-safes (greater than 5n+10), which indicate the absence of publication bias^62^. This meta-analysis, conducted with utmost care, showed no publication bias. The use of random-effects meta-analysis, a comprehensive approach, allowed us to assess the impact of microbial fertilizer applications on yield performance (*P*<0.0001).

## Results

### Reshaping root–microbiome symbiosis through increased nutrient supply

The application of organic fertilizer in addition to chemical fertilizer can enhance the availability of nutrients, such as SOC, AP, and AK, in soil without causing a significant change in soil pH (Table S1). The growth of wheat under different nutritional conditions is depicted in Fig. 1a. The plant height increased linearly from OF0 to OF20 and decreased at OF30. Treatments with high organic fertilizer application resulted in greater relative abundances of γ-*Proteobacteria*, *Agaricomycetes*, and *Orbiliomycetes* and lower relative abundances of *Bacilli* and *Sordariomycetes* (Fig. 1b). Unlike the bacterial richness (Shannon indice), which did not vary significantly among different treatments (Fig. 1c), the fungal richness was significantly greater under high levels of organic fertilizer application (Fig. 1d). During bacterial and fungal community assembly, stochastic processes dominated in all treatments and significantly decreased in the bacterial community under OF30 according to the normalized stochasticity ratio (Fig. S1a-b). Additionally, the wheat yield (Fig. 1e) and grain number (Fig. 1f) peaked under the OF20 treatment, while the average grain weight was greatest under the OF5 treatment (Fig. 1g). We conducted a partial Mantel test to examine the correlation between geographical distance-corrected dissimilarities of community composition and environmental variables (Fig. 1h). Our results showed that both the bacterial and fungal community compositions were significantly correlated with SOC, NO_3_^-^-N, and AK. We found that the bacterial community composition was correlated with grain number and that SOC, NO_3_^-^-N, AK, and AP were positively correlated with grain number. Notably, SOC was identified as the most significant factor influencing wheat yield (Fig. 1i). Moreover, the predicted microbial functions involved in nitrogen cycling increased with the application of organic fertilizer, showing a strong correlation with soil available nutrients, wheat yield and grain number (Fig. S1 c-d). Additionally, co-occurrence network analysis between different taxas and wheat yield revealed that the fungal class *Orbiliomycetes* played a crucial role in crop yield and grain number (Fig. 1j).

**Fig. 1.**
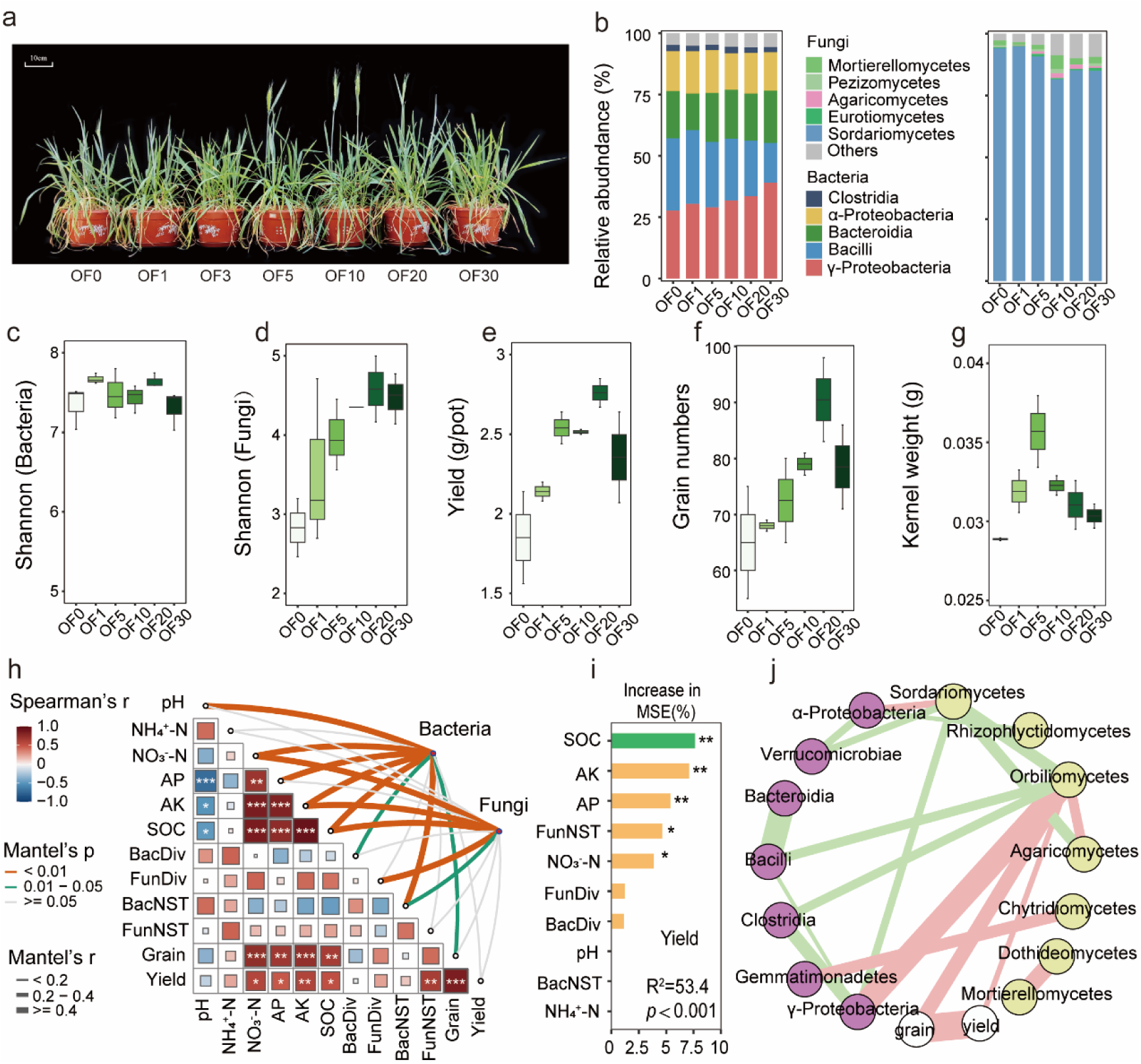
Reshaping root–microbiome symbiosis with increasing amounts of nutrients. **a,** Wheat grown with increasing amounts of organic fertilizer: 0% (OF0), 1% (OF1), 3% (OF3), 5% (OF5), 10% (OF10), 20% (OF20), and 30% (OF30). **b,** Changes in the rhizosphere microbial community under different organic fertilizer treatments. **c-d,** Bacterial (c) and fungal (d) richness (Shannon) among all soil samples. **e-g,** Yield (e), grain number (f) and average kernel weight (g) of wheat plants grown under different conditions. **h,** Correlations between environmental/microbial factors and crop performance and microbial communities. Edge width corresponds to the absolute value of the correlation coefficient as determined by linear mixed-effects models. The colors indicate correlation types. The solid and dashed lines denote significant and nonsignificant correlations, respectively, based on Wald type II χ² tests. Pairwise comparisons of environmental/microbial factors and crop performances are shown in the triangle. Spearman’s correlation coefficient is shown with a color gradient. **i,** Key factors affecting wheat yield according to random forest analysis. Bar charts indicate the relative importance of variables as predictors of wheat yield. **j,** Co-occurrence network between microbial class level and wheat yield and grain number.

### Nutrition-dependent fine-tuning of the root microbiome with different bioinoculant components

The microbial community composition of EMs, a widely used commercial bioinoculants worldwide, is depicted in Fig. 2a. The ASVs (Amplicon Sequence Variants) in EM that successfully colonized were mainly fungi, particularly *Eurotiomycetes* and *Saccharomycetes*, as indicated in the evolutionary tree of microbial species from EM (Fig. 2a-b). 6 fungal ASVs were found to successfully colonize the wheat rhizosphere after EM pellet application in OF0, while only one and none were able to colonize under OF5 and OF20 conditions, respectively. Following EM application, one ASV was able to colonize the wheat rhizosphere, and significant changes were detected in five ASVs after EM supernatant application (Fig. 2b-c). Interestingly, the colonized ASV numbers following pellet application exhibited a strong linear correlation with the nutrient level. The rhizosphere bacterial and fungal community composition (Fig. S2a-b), diversity (Fig. S2c-d) and normalized stochasticity ratio (Fig. S2e-f) after EM, EM supernatant and EM pellet application under the three fertilization conditions are depicted in Fig. S2. In the OF20 treatment, both the bacterial and fungal shannon indices as well as the NST index decreased after EM pellet application. Conversely, in the OF5 treatment, only the fungal shannon (Fig. S2c-d) and NST indices (Fig. S2e-f) decreased following pellet application.

**Fig. 2.**
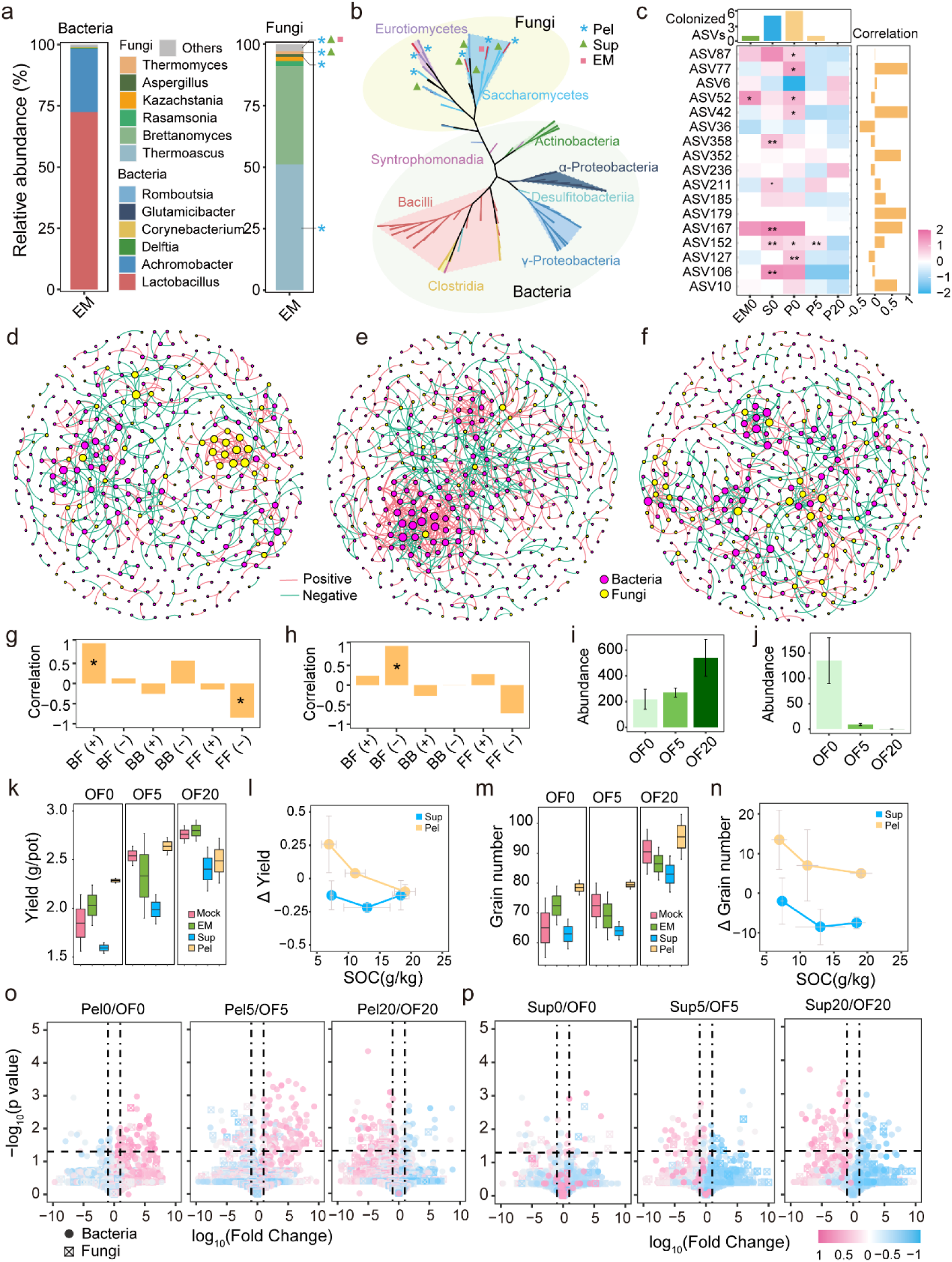
Nutrition-dependent fine-tuning of the root microbiome with different bioinoculant components. **a-b,** EM microbial community composition (a) and phylogenetic relationships between different microbial ASVs (b). Asterisks, triangles and quadrilaterals indicate ASVs successfully colonized in the root rhizosphere after pellet, supernatant and whole EM inoculant application, respectively. **c,** Heatmap and histogram showing absolute abundance fold changes in ASVs colonized in the rhizosphere (* indicates ASVs colonized under different EM components and OF applications). The histogram at the top displays the numbers of ASVs, while the histogram on the right shows the correlations between ASVs and the amount of OF applied. **d-f,** Co-occurrence network analysis of the full dataset (OF and EM application) showing that microbial interkingdom network patterns differed among the OF0 (d), OF5 (e), and OF20 (f) treatments. **g-h,** Histogram illustrating the correlations between microbial interaction lines and the amount of nutrient applied in the rhizosphere (g) or between EM and the rhizosphere (h). BF (+): positive interactions between bacteria and fungi; BF (-): negative interactions between bacteria and fungi; BB (+): positive interactions between bacteria; BB (-): negative interactions between bacteria; FF (+): positive interactions between fungi; FF (-): negative interactions between fungi. **i,** Changes in the absolute abundance of antagonistic bacteria when EM was applied under three different organic fertilizer levels. **j,** Changes in the absolute abundance of fungi colonizing the roots when EM was applied under three different organic fertilizer levels. **k-n,** Changes in the wheat yield (k) and grain number (m) with the addition of the EM, EM supernatant and EM pellets under different levels of organic fertilizer. Linear graphs showing the changes in yield (l) and grain number (n) after EM supernatant and EM pellet application. **o-p,** Volcano plots showing upregulated and downregulated bacterial and fungal ASVs after EM pellet (o) and EM supernatant (p) application. The dotted line on the x-axis is utilized to determine whether the ASVs in the EM pellet or EM supernatant treatment group increased or decreased compared to those in the CK group, with lower values on the left and higher values on the right. The gray line on the y-axis was determined by the value of -log_10_(0.05), which represents the log_10_ value of *p* = 0.05.

We conducted a co-occurrence network analysis to further evaluate the impact of EM pellet (Fig. 2d-f) or EM supernatant (Fig. S3a-c) application on bacterial-fungal interkingdom interactions in the wheat rhizosphere under the three OF treatments. We found that fungal modules exhibited greater network connectivity (i.e., network degree) than that in bacterial taxa under OF0 (Fig. 2d), but this trend was reversed under OF5 (Fig. 2e). In OF5, the degree of bacterial taxa is significantly correlated with each taxa’s grain number and yield after EM pellet application (Fig. S5k, n). Fungal taxa degree significantly correlated with each taxa’s yield in OF20 (Fig. S5l). No significant correlations were observed in other treatments (Fig. S3j-o). Furthermore, modules of bacterial ASVs with high node degrees after pellet application (Fig. S5a-b) demonstrated a strong positive correlation with wheat yield (Fig. S5d-e) and grain number (Fig. S5g-i). In contrast, most fungal ASVs showed a negative correlation with wheat yield (Fig. S5d-f) and grain number (Fig. S5g-i). After EM supernatant treatments, bacteria played an increasingly significant role with higher OF application in microbial network analysis (Fig. S3a-c) and modules of both bacterial and fungal ASVs did not exhibit strong correlations (Fig. S3d-i). Fungal taxa had higher network connectivity than bacterial taxa in OF0, while the pattern was reversed in OF20 (Fig. S3a-c). The distribution of grain number or yield-correlated ASVs is shown in Fig. S3d-i. High-degree bacterial and fungal taxa generally negatively correlated with grain numbers in OF0 and OF20 (Fig. S3d-f). In contrast, high-degree bacterial taxa positively correlated with yield in OF20 (Fig. S3g-i). In the OF0 treatment, bacterial taxa degree showed a significant negative correlation with yield relevance (Fig. S3j), while no significant correlations were found in other treatments (Fig. S3k-o).

We further isolated network edges between microbial ASVs detected in the EM and wheat rhizosphere in the networks after pellet application. The results indicated that positive interactions between bacterial and fungal ASVs in the rhizosphere, as well as negative interactions between fungal ASVs from EM and bacterial ASVs in the rhizosphere, were significantly correlated with the amount of nutrient application level (Fig. 2g-h). The absolute number of bacterial ASVs negatively correlated with colonized fungi from EM increased with OF application (Fig. 2i), and the changes in the absolute number of bacterial/fungal ASVs positively/negatively correlated with colonized fungi from EM are shown in Fig. S4b-d. The highest absolute number of fungal ASVs colonizing the rhizosphere was found in OF0 (Fig. 2j). We conducted correlation analysis between different types of edge numbers and the colonization efficiency of ASVs from EM (Fig. S4a). The results revealed positive interactions between bacterial and fungal ASVs detected only in the rhizosphere, as well as negative interactions between ASVs detected in the EM and rhizosphere that played active roles in EM colonization. In contrast, negative interactions between fungal ASVs in the rhizosphere showed a strong positive correlation with colonization efficiency. The number of positive and negative edges between bacterial ASVs in the rhizosphere and fungal ASVs from the EM increased under higher OF applications, while the opposite trend was found for the number of negative interaction lines between fungal ASVs from the EM and the rhizosphere (Fig. S4e-g). Under the three different fertilization treatments, the wheat yield decreased after supernatant application. The yield increased after pellet application under 0% and 5% organic fertilizer application but decreased when 20% organic fertilizer was applied (Fig. 2k). The wheat yield response to pellet application decreased with increasing soil organic carbon (SOC) concentration (Fig. 2l). The grain number decreased after supernatant application but increased after pellet application in all three treatments (Fig. 2m). The response of the grain number to pellet application also decreased with the SOC concentration (Fig. 2n).

We compared microbial community changes after EM pellet (Fig. 2o) and EM supernatant (Fig. 2p) application under three different conditions. Under OF0 and OF5 fertilization conditions, pellet treatment led to an increase in grain number, and 48 of 51 ASVs in OF0 and 103 of 139 ASVs in OF5, which were positively correlated with grain yield, were upregulated. 11 of the 12 ASVs in OF0 and 34 of the 44 ASVs in OF5, which were negatively correlated with grain yield, were downregulated. Under OF20 conditions, pellet treatment led to an increase in grain number and a decrease in yield. 72 of the 73 ASVs that were positively correlated with grain yield were also downregulated (Fig. 2o & Table S2). In contrast, supernatant treatment resulted in a decrease in grain number, and 4 of the 11 ASVs in OF0, 16 of the 17 ASVs in OF5, and 38 of the 40 ASVs in OF20, which were negatively correlated with grain production, were upregulated. 11 of the 19 ASVs in OF0, 25 of the 27 ASVs in OF5, and 75 of the 76 ASVs in OF20 were positively correlated and were downregulated (Fig. 2p & Table S2). With increasing organic fertilizer application, more fungal and bacterial ASVs, which were positively correlated with wheat yield, were downregulated with the addition of the supernatant (Fig. S6a-d). Fungal ASVs, which are negatively correlated with grain number, were downregulated with pellet application. Bacterial ASVs, which were correlated with grain number, were downregulated with supernatant application (Fig. S7a-d).

### The effect of microbe and microbial functional coordination on wheat yield and the root-associated microbiome under EM pellet application

Compared with Firmicutes, the bacterial phyla Bacteroidetes and Proteobacteria were predominantly enriched after pellet application (Fig. 3a-c). In the OF5 treatment, most ASVs showed significant enrichment or depletion, whereas, the fewest ASVs exhibited significant changes under the OF0 treatment. In the OF20 treatment, much fewer ASVs from *Proteobacteria* were unregulated than in the other treatment groups. Additionally, we analyzed the changes in FAPROTAX functions after pellet application. Nitrite respiration- and denitrification-related functions were significantly enriched after pellet application under OF0 (Fig. 3g) or OF5 (Fig. 3h) conditions but were significantly depleted under conditions with OF20 application (Fig. 3i). Furthermore, we examined the functions of ASVs that were positively correlated with crop yield. Nitrogen cycling-related functions, which were enriched under low OF application conditions and depleted under conditions with high OF application, were found to be involved in regulating crop yield. Random forest analysis revealed that nitrite respiration was the most crucial function correlated with crop yield (Fig. 3j). After supernatant application, *Proteobacteria* was significantly depleted in the rhizosphere of OF20 treatment (Down: 44, Up: 14). However, other bacterial phyla, including *Firmicutes* and *Bacteroidota*, showed no significant difference in all treatments (Fig. S8a-c). ASVs of Fungal phyla Ascomycota exhibited the most downregulations under OF20 (Fig. S8d-f). Microbial functions, such as photoheterotrophy and phototrophy, were significantly downregulated under OF0 (Fig. S8g), while functions of cellulolysis and nitrogen fixation were significantly upregulated under OF5 (Fig. S8h). Furthermore, nitrogen conversion-related functions, including nitrate reduction and nitrate respiration, *etc*. were significantly down-regulated under OF20 (Fig. S8i). Also, nitrite denitrification and denitrification were determined to be the most crucial functions correlated with crop yield performance (Fig S8j).

**Fig. 3.**
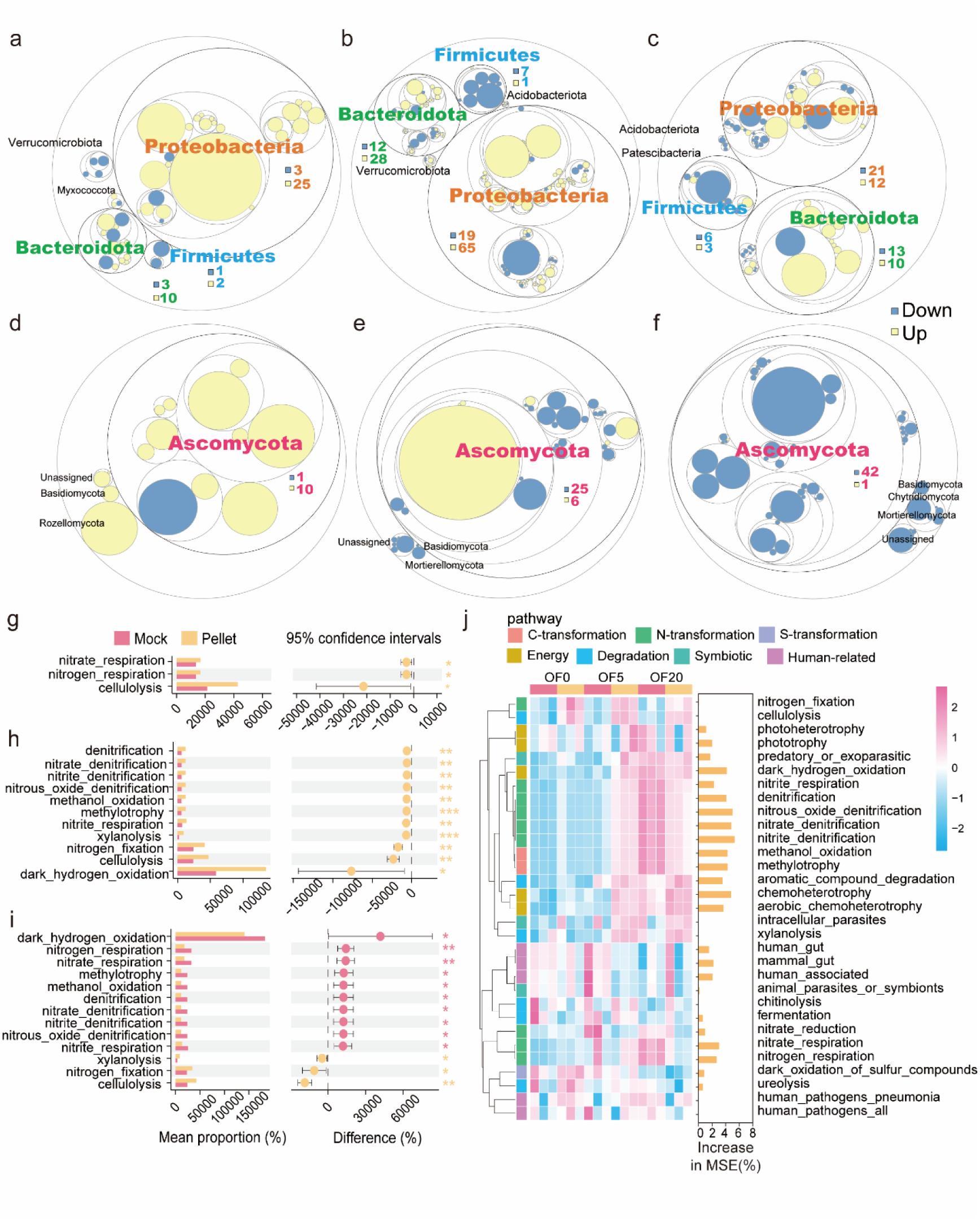
Functional coordination between wheat yield and the root-associated microbiome under EM pellet application. **a-f,** Differences in amplicon sequence variant (ASV) levels are depicted in the figure. The blue and yellow bubbles represent bacterial ASVs in OF0 (a), OF5 (b), and OF20 (c) and fungal ASVs in OF0 (d), OF5 (e), and OF20 (f) with significant enrichment (FDR < 0.05) in soil after pellet application, based on statistical analysis with ALDEx2. The size of the bubbles represents the effect size between the relative abundance of ASVs in the CK and EM pellet treatments as determined by ALDEx2. **g-i,** Changes in FAPROTAX functions following the application of EM pellets are depicted in the bar charts, which show the mean value and standard errors for each category under OF0 (g), OF5 (h) and OF20 (i). The dots on the chart represent the percentage change in effect size between the CK and EM pellets at 95% confidence intervals (CIs). **j,** Heatmap illustrating changes in bacterial FAPROTAX functions following the application of EM pellets, accompanied by a bar chart demonstrating the significance of these functions in influencing crop yield based on random forest analysis.

### Mechanisms of plant‒root microbiome symbiotic regulation under different nutritional conditions

To better distinguish the direct and indirect effects of environmental factors on microbial biodiversity after the application of the EM supernatant and pellet, we conducted structural equation modeling (SEM) analyses. Overall, wheat yield and grain number were enhanced directly by pellet application and suppressed by supernatant application under OF0, a condition without organic fertilizer application (Fig. 4a; Fig. S9a). This direct increase in wheat yield and grain number decreased with increasing organic fertilizer application (Fig. 4a-c). The supernatant had a direct inhibitory effect on wheat yield under all fertilization conditions (Fig. S9a-c). Furthermore, rhizosphere bacterial and fungal communities were found to play a more important role in regulating grain number and yield under conditions with greater organic fertilizer application. Under conditions involving OF5 and OF20 application, both the pellet and supernatant indirectly regulated the wheat yield and grain number by affecting microbial diversity and community assembly (Fig. 4b-c; Fig. S9b-c).

**Fig. 4.**
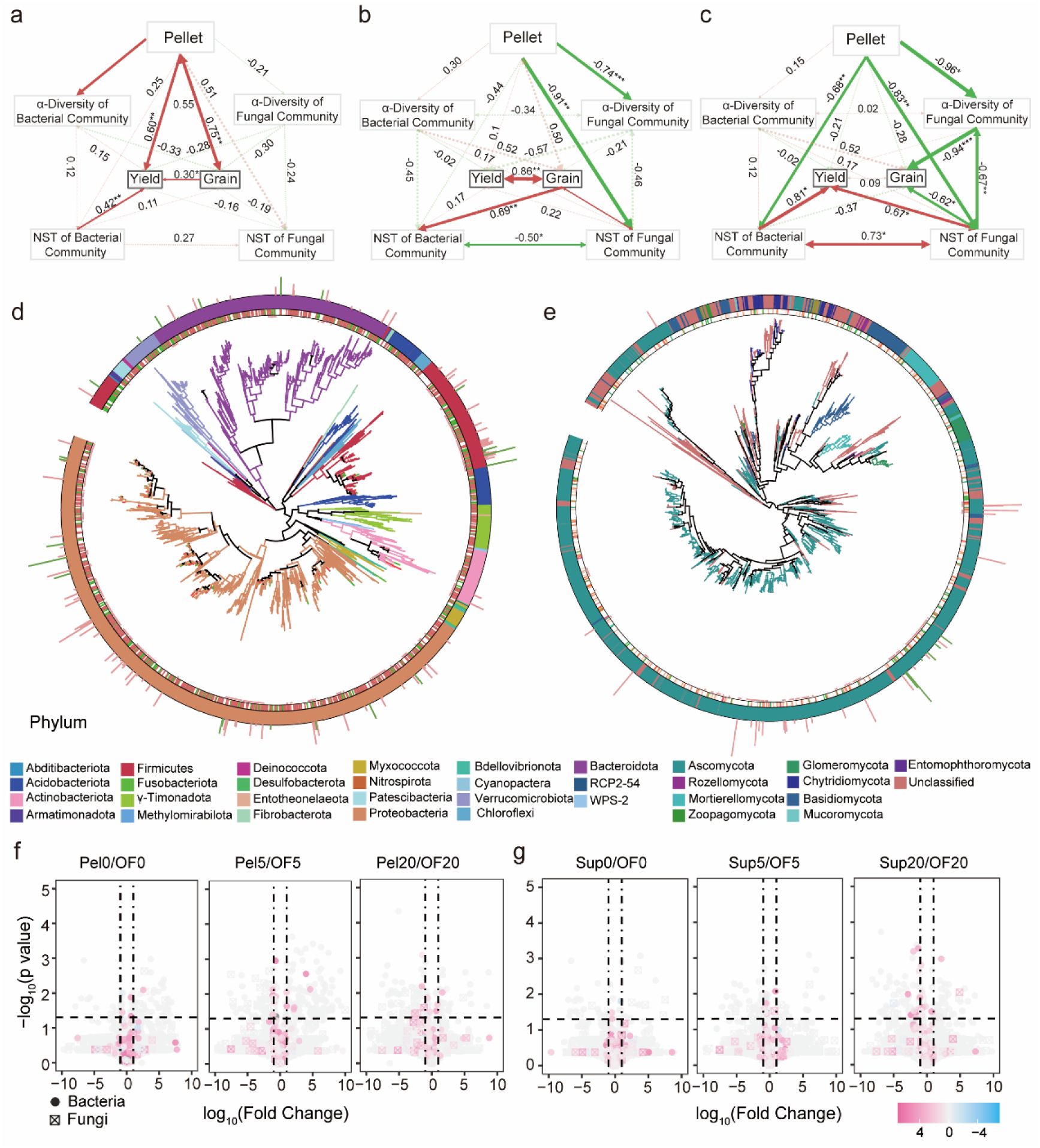
Mechanisms of plant ‒ root microbiome symbiosis regulation under different nutritional conditions. **a-c,** Structural equation models (SEMs) illustrating the relationships among treatments, plant variables, bacterial and fungal richness and NST. Panel (a) represents 0% OF; panel (b) represents 5% OF; and panel (c) represents 20% OF along with pellet application. Solid or dashed lines indicate significant (*p* < 0.05) or nonsignificant relationships, respectively. The numbers near the pathway arrow indicate the standard path coefficients. **d-e,** Phylogenetic trees based on 16S rRNA genes (d) and ITS genes (e) from root microbiome high-throughput sequencing illustrating the branching patterns that represent the common ancestry and evolutionary divergence of species. The branches of the tree depict evolutionary paths, with longer branches indicating greater divergence and shorter branches indicating closer evolutionary relationships. The innermost signs indicate ASVs isolated from wheat root microbiota using high-throughput cultivation technology. The first ring shows the correlations between ASVs and wheat yield, with red indicating positive correlations and green indicating negative correlations. The second ring displays the phylum levels of the ASVs. The outermost bars represent the effect of each ASV on wheat growth in terms of weight, with red indicating positive effects and green indicating negative effects. **f-g,** Volcano plots displaying the upregulation and downregulation of bacterial and fungal ASVs following the application of the EM pellets (f) and EM supernatant (g).

We isolated rhizosphere bacteria and fungi from wheat plants grown under pot conditions via previously established high-throughput cultivation and identification methods. A total of 2,073 wells containing a diverse range of bacterial isolates belonging to Proteobacteria, Actinobacteria, Firmicutes, and Bacteroidetes were successfully cultivated. These cultivated bacteria, with a total of 385 distinct ASVs, accounted for 66.6% of the wheat root-associated ASVs (Fig. 4e). Additionally, a total of 2073 wells containing 178 distinct ASVs of fungal isolates belonging to Ascomycota were cultivated successfully (Fig. 4f). These cultivated fungi accounted for 45.8% of the wheat root-associated fungal ASVs. Most cultivated bacteria and fungi were able to promote wheat growth (Fig. 4e-f). We determined the distribution of cultivable bacteria in the wheat rhizosphere through sequence alignment. In the OF0 and OF5 treatments, the EM pellets significantly upregulated bacterial ASV11 (*Rheinheimera*), ASV1 (*Pseudomonas*), and ASV49 (*Hydrogenophaga*), which had significant growth-promoting effects on wheat growth (Fig. 4g). However, no significant upregulation of growth-promoting microbes was found under conditions with supernatant application. However, four growth-promoting bacteria were downregulated under OF20 application after supernatant treatment (Fig. 4h). We compared the impact of cultured bacteria on wheat growth under three conditions: OF0, OF0 + EM pellet, and OF0 + EM supernatant. Under OF0, 147 growth-promoting and 19 growth-inhibiting bacterial ASVs were identified (Fig. S10a). Following the application of EM pellet, no significant difference was observed. 119 bacterial ASVs were found to promote wheat growth, while 17 bacteria ASVs exhibited an inhibitory effect (Fig. S10b). However, the application of supernatant led to a significant down-regulation in growth-promoting ASVs and an up-regulation in those with a growth inhibitory effect (Fig. S10c).

### Projected changes in wheat yield after the application of bioinoculants worldwide

The distribution of soil organic carbon (SOC) in the world’s wheat-growing areas exhibits a distinct spatial pattern, with higher values in countries at high latitudes, such as Russia, Canada, and Finland, and lower values in Africa and South America (Fig. 5a). Similar patterns were also found in China, with higher values in the southeastern Qinghai-Tibetan Plateau and northeastern China and lower values in the arid regions of northwestern China, where deserts and sandy lands are more common (Fig. S11a). We predicted the potential effects of EM (Fig. 5b), EM pellet (Fig. 5c) or EM supernatant (Fig. 5d) application on wheat yield based on the global distribution of SOC (Fig. S11b-e). As illustrated in Fig. 5b-e and Fig. S11b-e, wheat yield showed significant spatial heterogeneity after EM, EM pellet and EM supernatant treatments. The overall trend aligns with the mean value distributions. Over all, application of commercial EM led to a reduction in global wheat yield by 4.1%, while EM pellet results in a significant increase in global wheat yield by 17.3 % and EM supernatant led to a substantial decrease in global wheat yield by 16.6% (Fig. 5f). Pellet application typically enhanced wheat yield, particularly in Africa, the Middle East, and Australia, while reducing yields in Russia and Canada. Supernatant application decreases wheat yield globally, while EM application reduces yields across most regions of the world. We outlined the potential effects in China (Fig. S11b-f) and in the 9 other top wheat-producing countries (Fig. 5b-f). Pellet application generally leads to an increase in wheat yield in most countries, except in Russia, Australia, Ukraine, and Germany. Conversely, supernatant application tends to decrease wheat yield in all top-ranking wheat-producing countries. In India, pellet application can boost wheat yield by 29.1 % (Fig. 5e). In China, commercial EM led to a reduction of wheat yield by 3.5 %, EM pellet increased wheat yield by 12.7 % and EM supernatant reduced wheat yield by 16.8 % (Fig. S11g). EM pellet application can increase wheat yield in the main wheat producing areas. In Shandong Province, pellet application can increase wheat yield by 23.51% (Fig. S11f). And, EM supernatant led to a substantial decrease in wheat yield by 16.81% in China (Fig. S11g). To further confirm our prediction, a global meta-analysis proved bio-inoculants’ positive effect on most continents, with the highest in West Asia and Africa and the lowest in Oceania and North America (Fig. 5g). This trend was similar to the projected effect of EM pellet application on global wheat production. With the increase of SOC content, the positive effect of bio-inoculants on crop yield gradually diminishes. When the SOC content exceeds 35 mg/kg, bio-inoculants negatively impact crop yield (Fig. 5h-i).

**Fig. 5.**
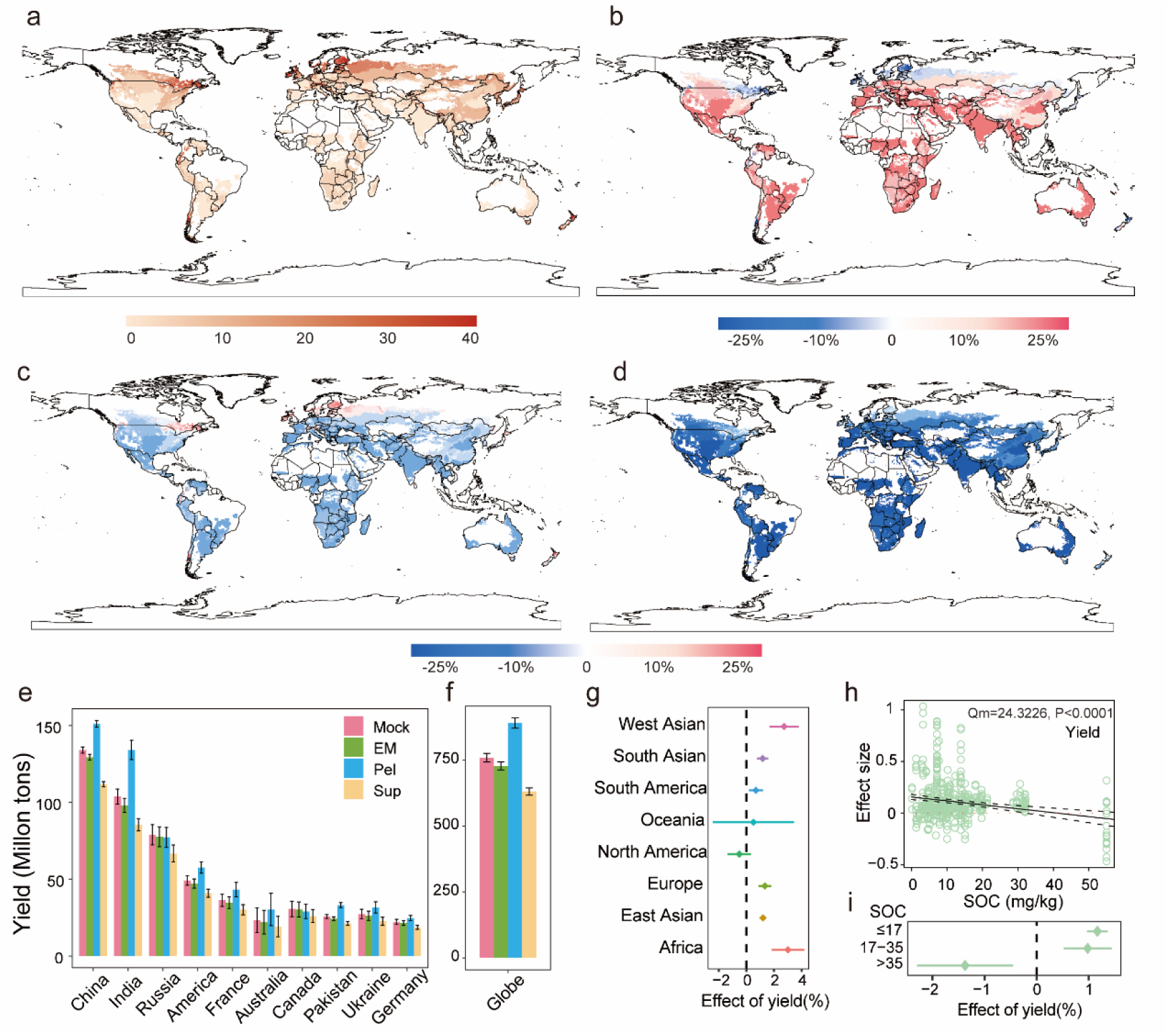
Global overview of changes in wheat yield after bioinoculant application. **a,** Spatial distribution of the mean SOC (kg C m^-2^) in the top 0-20 cm soil layer. **b-d,** Projected spatial effect of the EM (b), EM pellets (c) and supernatant (d) on the global wheat yield. **e,** Predicted effects of EM and its components on wheat yield in the top 10 wheat-producing countries over the past 5 years. **f,** Projected effect of EM and its pellet or supernatant on wheat yield in the world. **g**, Effect size of microbial fertilizer on crop yield in different regions of the world. **h-i**, Effect size of soil organic carbon (SOC) content on crop yield globally.

**Fig. 6.**
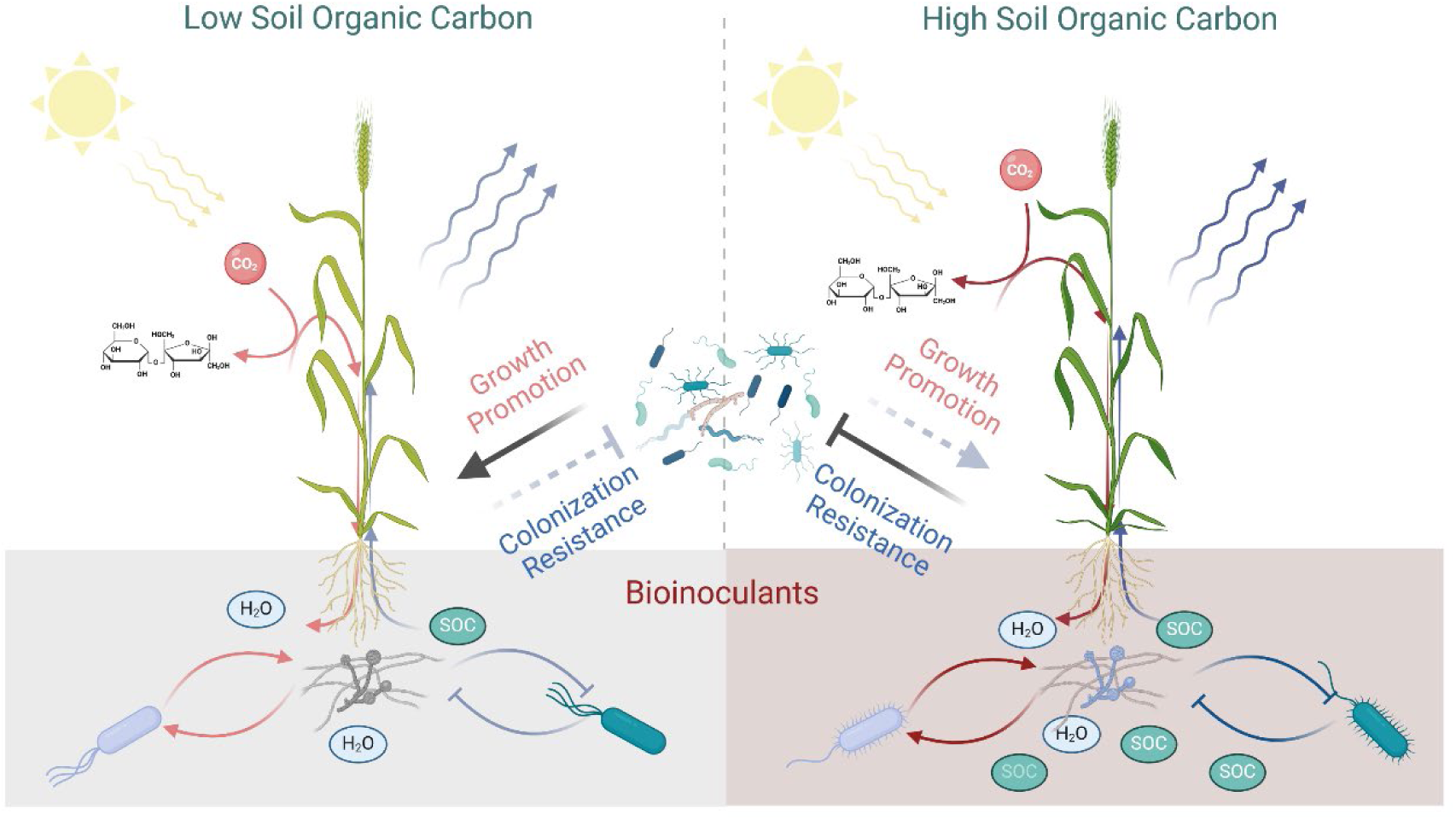
Schematic diagram of nutrition-dependent wheat yield regulation by live microbial inoculants. As the most crucial factor in promoting yield in commercial EM, EM pellets directly regulate wheat yield by increasing the number of wheat grains. Soil organic carbon (SOC) is the key soil property that shapes the interaction of soil microbial communities by enhancing the positively correlated interactions between rhizosphere bacteria and fungi. These interactions in the rhizosphere play a important role in preventing bioinoculant invasion by antagonizing bacteria. Therefore, under conditions with low SOC content, EM pellets were found to enhance wheat yield. However, due to colonization resistance and the trade-off between grain number and grain weight, application of EM pellets under conditions of high SOC content led to a decrease in yield.

## Discussion

Soil microorganisms are abundant and resource-rich and play a significant role in activating nutrients and enhancing crop resistance to biotic or abiotic stresses. They have been widely utilized as bioinoculants, biocontrol agents, or biostimulants^7,10,27^. However, our understanding of the process leading to successful colonization and persistence in plant microbial communities is still limited, and transitioning beneficial microorganisms from laboratory or greenhouse settings to field conditions for regulating plant phenotypes is challenging due to the influence of climate, the soil environment, crop genotype, and other factors. Although many microbial strains demonstrate promising plant productivity characteristics under experimental conditions, their application in the field has often been unsuccessful^13,28^. Several studies have reported that the abundance and activity of arbuscular mycorrhizal fungi (AMF) can be explained by the soil microbiome and by pesticide application^29-31^. Moreover, the favorability of laboratory settings for the growth of microbes may contribute to the potentially robust colonization of crop root microbial communities by bioinoculants in the laboratory^32^. This challenge is not unique to agriculture but also extends to clinical practice. For instance, during fecal microbiota transplantation (FMT), the success rate can be affected by host genotype, immune status, and intestinal microbial composition^14,33-36^.

Effective microorganisms (EMs) are mixtures of beneficial, naturally occurring microorganisms that can be used to increase the microbial diversity of soil ecosystems^37^. To date, EMs have become one of the most successful commercial bioinoculants used in a wide range of fields. Similar to the effect of other bioinoculants, that of EMs on soil productivity remains under debate^38,39^. Our results indicated that the effect of EMs arises from a combination of the growth-promoting effect of the EM pellets and the inhibitory effect of the EM supernatant. To our surprise, none of the bacterial species isolated from the EM pellets were able to colonize the wheat rhizosphere, and only a few fungal species could successfully colonize the wheat rhizosphere. In addition, the inhibitory effect of the EM supernatant did not depend on the soil nutrient status. We found that commercial EM application would lead to a reduction in wheat yield of 4.1% worldwide and 3.5% in China, and it would increase wheat yield only in northern Europe and the central region of North America. However, the use of cell pellets from EM could lead to a significant increase in global wheat yield by 17.3% and by 12.7% in China. Moreover, the EM supernatant would have a substantial inhibitory effect on global wheat production, with reductions of 16.6% worldwide and 16.8% in China (Fig. 5f; Fig. S11g). All these results indicated that even though the cell pellets in EM can increase crop yield performance, there are some unknown chemicals that might inhibit grain number. In agricultural practice, applying EM pellets or eliminating the inhibitory effects of EM supernatants has the potential to further improve the stability or yield-promoting effect of current commercial EM products. In addition, it will be important to identify chemicals and related chemical-producing bacteria or fungi arising from EM products in the future.

Nutrients are substances that provide the energy and biomolecules necessary for the proper functioning and growth of all living organisms. In nature, nutrient availability represents one of the most important environmental cues for shaping global cellular resource investment^40,41^, the strength of species interactions, the stability of microbial communities and multitrophic interactions in different ecosystems^19^. Elevated soil base nutrients (NH_4_^+^-N, NO_3_^−^-N, available P and available N) can amplify the effect of microbial inoculants on soil microbial biomass, such as fungal biomass, actinobacterial biomass, microbial biomass carbon, and microbial biomass nitrogen^42^. SOC represents the amount of carbon that is retained in the soil after the decomposition of organic matter^43-45^. It is a vital indicator that determines soil quality, fertility, agricultural profitability, and atmospheric carbon dioxide (CO_2_) fixation. As a nutrient reservoir, SOC provides a steady supply of essential elements for plant uptake and microbial growth in the soil. It has long been hypothesized that higher SOC contents can increase crop yield in food production systems^46-48^. However, to date, the relationship between nutrient availability and the interaction between bioinoculants and the crop root microbiome remains largely unclear.

In this study, we used a wheat production system as an example, supplemented the soil nutrient supply with different amounts of peach organic fertilizer, and investigated the interactions between different components of commercial bioinoculants and root–microbiome symbiosis in wheat. We initially conducted a comprehensive examination of crop growth and the rhizosphere microbial community under various nutrient conditions by providing different amounts of organic fertilizers in a pot experiment. Interestingly, the soil nutrient supply, fungal community assembly and grain number were found to be essential for regulating wheat yield, and SOC was found to be the most important factor. EM pellets increased grain number in a nutrient-dependent manner, while the EM supernatant had an inhibitory effect on grain number. However, under high organic fertilizer application conditions, EM pellets had a negative effect on wheat yield even when the grain number increased. This bottleneck was mainly caused by the tradeoff between average grain weight and grain number^49,50^. In the future, molecular breeding approaches could be used to overcome this bottleneck^51^.

Structural equation analysis revealed that the EM pellets were able to increase grain number and crop yield in a direct manner under no organic fertilizer supply. However, under higher organic fertilizer supply conditions, the contribution of the direct yield-promoting effect decreased, and the indirect inhibitory effect mediated by root fungal communities became more important in regulating grain number and crop yield (Fig. 4a-c). Moreover, successful colonization of bioinoculants in the crop rhizosphere was regulated in a nutrient-dependent manner by positive correlations between rhizosphere bacteria and fungi (Fig. 2g, i). These interactions prevented the colonization of bioinoculants by antagonizing bacteria, which can also be stimulated by a higher organic fertilizer supply (Fig. 2 g, h).

In conclusion, our results indicated that crop–root microbiome symbiosis can be reshaped by adjusting the soil organic carbon concentration. Bacterial and fungal interactions, most likely cooperative ones, in the rhizosphere were sensitive to soil organic carbon content, and they might have played an important role in preventing bioinoculant from colonizing the rhizosphere by antagonizing bacteria. Global meta-analysis indicated that these nutrient-dependent crop yield regulation mechanisms operate not only for EM pellets but also for other microbial inoculants. Soil organic carbon is possibly one of the most important nutritional determinants that shapes the phenotypic plasticity mediated by bioinoculants in the regulation of wheat yield. However, soil organic carbon is vulnerable to several global change stressors, including climate (drought, heatwaves, *etc*.) and anthropogenic (land-use intensification) factors. An increase in the number of global change stressors can decrease soil carbon storage and persistence across terrestrial ecosystems^52^. According to our result, using microbial inoculants under these circumstances will have the potential to enhance the promotion of global crop yield and overcome the stressors of global climate change on soil quality.

## Acknowledgments

This research was supported by the National Key R&D Program of China (2023YFD1900504, 2022YFD1901304), the National Natural Science Foundation of China (32100095) and the Chinese Universities Scientific Fund (2022RC033, 2023RC016, 2024RC025) to B.N.

## Author Contribution Statement

B.N. designed the research project; Y.H., S.T., J.L., and T.X. performed the experiments; Y.H. and S.T. wrote the first manuscript; B.N. edited the manuscript; and all the authors read and approved the final manuscript.

## Competing interests Statement

The authors declare no competing interests.

## Data availability

Bacterial 16S and fungal ITS data for wheat rhizosphere microbial community analysis in this paper were deposited in the Sequence Read Archive (http://www.ncbi.nlm.nih.gov/sra) under the BioProject ID PRJNA1124597 and PRNA1124615.

## Notes

### Competing Interest Statement

The authors have declared no competing interest.

## References

1 Tilman, D., Balzer, C., Hill, J. & Befort, B. L. Global food demand and the sustainable intensification of agriculture. Proc Natl Acad Sci U S A 108, 20260–20264 (2011).

2 Molotoks, A. et al. Global projections of future cropland expansion to 2050 and direct impacts on biodiversity and carbon storage. Glob Chang Biol 24, 5895–5908 (2018).

3 Anstalt, S. V. Food and agriculture organization of the United Nations. (2013).

4 Li, A. et al. Wheat breeding history reveals synergistic selection of pleiotropic genomic sites for plant architecture and grain yield. Mol Plant 15, 504–519 (2022).

5 Brenchley, R. et al. Analysis of the bread wheat genome using whole-genome shotgun sequencing. Nature 491, 705–710 (2012).

6 Peng, J. et al. ’Green revolution’ genes encode mutant gibberellin response modulators. Nature 400, 256–261 (1999).

7 Trivedi, P., Leach, J. E., Tringe, S. G., Sa, T. & Singh, B. K. Plant-microbiome interactions: from community assembly to plant health. Nat Rev Microbiol 18, 607–621 (2020).

8 Finkel, O. M. et al. A single bacterial genus maintains root growth in a complex microbiome. Nature 587, 103–108 (2020).

9 Duran, P. et al. Microbial Interkingdom interactions in roots promote *Arabidopsis* survival. Cell 175, 973–983 e914 (2018).

10 Singh, B. K., Trivedi, P., Egidi, E., Macdonald, C. A. & Delgado-Baquerizo, M. Crop microbiome and sustainable agriculture. Nat Rev Microbiol 18, 601–602 (2020).

11 Albright, M. B. N. et al. Solutions in microbiome engineering: prioritizing barriers to organism establishment. ISME J 16, 331–338 (2022).

12 Lawson, C. E. et al. Common principles and best practices for engineering microbiomes. Nat Rev Microbiol 17, 725–741 (2019).

13 Poppeliers, S. W., Sanchez-Gil, J. J. & de Jonge, R. Microbes to support plant health: understanding bioinoculant success in complex conditions. Curr Opin Microbiol 73, 102286 (2023).

14 Porcari, S. et al. Key determinants of success in fecal microbiota transplantation: From microbiome to clinic. Cell Host Microbe 31, 712–733 (2023).

15 Philippot, L., Raaijmakers, J. M., Lemanceau, P. & van der Putten, W. H. Going back to the roots: the microbial ecology of the rhizosphere. Nat Rev Microbiol 11, 789–799 (2013).

16 Rodriguez, P. A. et al. Systems biology of plant-microbiome interactions. Mol Plant 12, 804–821 (2019).

17 Dai, T. et al. Nutrient supply controls the linkage between species abundance and ecological interactions in marine bacterial communities. Nature communications 13, 175 (2022).

18 Ratzke, C., Barrere, J. & Gore, J. Strength of species interactions determines biodiversity and stability in microbial communities. Nat Ecol Evol 4, 376–383 (2020).

19 Zhu, L. et al. Resource-dependent biodiversity and potential multi-trophic interactions determine belowground functional trait stability. Microbiome 11, 95 (2023).

20 Kedia, S. et al. Faecal microbiota transplantation with anti-inflammatory diet (FMT-AID) followed by anti-inflammatory diet alone is effective in inducing and maintaining remission over 1 year in mild to moderate ulcerative colitis: a randomised controlled trial. Gut 71, 2401–2413 (2022).

21 Jang, Y. O. et al. Fecal microbial transplantation and a high fiber diet attenuates emphysema development by suppressing inflammation and apoptosis. Exp Mol Med 52, 1128–1139 (2020).

22 Lal, R. Soil carbon sequestration impacts on global climate change and food security. Science 304, 1623–1627 (2004).

23 Wardle, D. A. et al. Ecological linkages between aboveground and belowground biota. Science 304, 1629–1633 (2004).

24 Zhao, S. et al. A precision compost strategy aligning composts and application methods with target crops and growth environments can increase global food production. Nat Food 3, 741–752 (2022).

25 Ikoyi, I. et al. Responses of soil microbiota and nematodes to application of organic and inorganic fertilizers in grassland columns. Biology and Fertility of Soils 56, 647–662 (2020).

26 Pergola, M. et al. Composting: The way for a sustainable agriculture. Applied Soil Ecology 123, 744–750 (2018).

27 Jansson, J. K., McClure, R. & Egbert, R. G. Soil microbiome engineering for sustainability in a changing environment. Nat Biotechnol 41, 1716–1728 (2023).

28 Russ, D., Fitzpatrick, C. R., Teixeira, P. & Dangl, J. L. Deep discovery informs difficult deployment in plant microbiome science. Cell 186, 4496–4513 (2023).

29 Lutz, S. et al. Soil microbiome indicators can predict crop growth response to large-scale inoculation with arbuscular mycorrhizal fungi. Nat Microbiol 8, 2277–2289 (2023).

30 Svenningsen, N. B. et al. Suppression of the activity of arbuscular mycorrhizal fungi by the soil microbiota. ISME J 12, 1296–1307 (2018).

31 Edlinger, A. et al. Agricultural management and pesticide use reduce the functioning of beneficial plant symbionts. Nature Ecology & Evolution 6, 1145–1154 (2022).

32 Jiang, M. et al. Home-based microbial solution to boost crop growth in low-fertility soil. New Phytol 239, 752–765 (2023).

33 Wu, L. et al. Data-driven prediction of colonization outcomes for complex microbial communities. Nat Commun 15, 2406 (2024).

34 Xiao, Y., Angulo, M. T., Lao, S., Weiss, S. T. & Liu, Y. Y. An ecological framework to understand the efficacy of fecal microbiota transplantation. Nat Commun 11, 3329 (2020).

35 Caballero-Flores, G., Pickard, J. M. & Nunez, G. Microbiota-mediated colonization resistance: mechanisms and regulation. Nat Rev Microbiol 21, 347–360 (2023).

36 Aggarwala, V. et al. Precise quantification of bacterial strains after fecal microbiota transplantation delineates long-term engraftment and explains outcomes. Nat Microbiol 6, 1309–1318 (2021)

37 Higa, T. Effective microorganisms: a biotechnology for mankind (2003)

38 Mayer, J., Scheid, S., Widmer, F., Fliessbach, A. & Oberholzer, H. R. How effective are ‘Effective microorganisms® (EM)’? Results from a field study in temperate climate. Applied Soil Ecology 46, 230–239 (2010).

39 Talaat, N. B. Effective microorganisms: An innovative tool for inducing common bean (*Phaseolus vulgaris* L.) salt-tolerance by regulating photosynthetic rate and endogenous phytohormones production. Sci Hortic-Amsterdam 250, 254–265 (2019).

40 Klumpp, S., Zhang, Z. & Hwa, T. Growth rate-dependent global effects on gene expression in bacteria. Cell 139, 1366–1375 (2009).

41 Ni, B., Colin, R., Link, H., Endres, R. G. & Sourjik, V. Growth-rate dependent resource investment in bacterial motile behavior quantitatively follows potential benefit of chemotaxis. Proc Natl Acad Sci U S A 117, 595–601 (2020).

42 Li, C. et al. Meta-analysis reveals the effects of microbial inoculants on the biomass and diversity of soil microbial communities. Nat Ecol Evol (2024).

43 Tao, F. et al. Microbial carbon use efficiency promotes global soil carbon storage. Nature 618, 981–985 (2023).

44 Schmidt, M. W. I. et al. Persistence of soil organic matter as an ecosystem property. Nature 478, 49–56 (2011).

45 Wieder, W. R., Bonan, G. B. & Allison, S. D. Global soil carbon projections are improved by modelling microbial processes. Nat Clim Change 3, 909–912 (2013).

46 Oldfield, E. E., Bradford, M. A. & Wood, S. A. Global meta-analysis of the relationship between soil organic matter and crop yields. Soil 5, 15–32 (2019).

47 Ma, Y. et al. Global crop production increase by soil organic carbon. Nature Geoscience 16, 1159–1165 (2023).

48 Lal, R. Soil carbon sequestration impacts on global climate change and food security. Science 304, 1623–1627 (2004).

49 Xie, Q. & Sparkes, D. L. Dissecting the trade-off of grain number and size in wheat. Planta 254 (2021).

50 Quintero, A., Molero, G., Reynolds, M. P. & Calderini, D. F. Trade-off between grain weight and grain number in wheat depends on GxE interaction: A case study of an elite CIMMYT panel (CIMCOG). Eur J Agron 92, 17–29 (2018).

51 Calderini, D. F. et al. Overcoming the trade-off between grain weight and number in wheat by the ectopic expression of expansin in developing seeds leads to increased yield potential. New Phytol 230, 629–640 (2021).

52 Saez-Sandino, T. et al. Increasing numbers of global change stressors reduce soil carbon worldwide. Nat Clim Change (2024).

53 Bai, Y. et al. Functional overlap of the *Arabidopsis* leaf and root microbiota. Nature 528, 364–369 (2015).

54 Zhang, J. et al. High-throughput cultivation and identification of bacteria from the plant root microbiota. Nat Protoc 16, 988–1012 (2021).

55 Magoc, T. & Salzberg, S. L. FLASH: fast length adjustment of short reads to improve genome assemblies. Bioinformatics 27, 2957–2963 (2011).

56 Bokulich, N. A. et al. Quality-filtering vastly improves diversity estimates from Illumina amplicon sequencing. Nat Methods 10, 57–59 (2013).

57 Edgar, R. C., Haas, B. J., Clemente, J. C., Quince, C. & Knight, R. UCHIME improves sensitivity and speed of chimera detection. Bioinformatics 27, 2194–2200 (2011).

58 Haas, B. J. et al. Chimeric 16S rRNA sequence formation and detection in Sanger and 454-pyrosequenced PCR amplicons. Genome Res 21, 494–504 (2011).

59 Ning, D. L., Deng, Y., Tiedje, J. M. & Zhou, J. Z. A general framework for quantitatively assessing ecological stochasticity. P Natl Acad Sci USA 116, 16892–16898 (2019).

60 Terrer, C., Vicca, S., Hungate, B. A., Phillips, R. P. & Prentice, I. C. Mycorrhizal association as a primary control of the CO_2_ fertilization effect. Science 353, 72–74 (2016).

61 Li, L. D., Hong, M., Zhang, Y. & Paustian, K. Soil N_2_O emissions from specialty crop systems: A global estimation and meta-analysis Global Change Biol 30 (2024).

62 Yao, Z. S., et al. A global meta-analysis of yield-scaled N_2_O emissions and its mitigation efforts for maize, wheat, and rice Global Change Biol 30 (2024).

